# HELLO: A hybrid variant calling approach

**DOI:** 10.1101/2020.03.23.004473

**Authors:** Anand Ramachandran, Steven S. Lumetta, Eric Klee, Deming Chen

## Abstract

Next Generation Sequencing (NGS) technologies that cost-effectively characterize genomic regions and identify sequence variations using short reads are the current standard for genome sequencing. However, calling small indels in low-complexity regions of the genome using NGS is challenging. Recent advances in Third Generation Sequencing (TGS) provide long reads, which call large-structural variants accurately. However, these reads have context-dependent indel errors in low-complexity regions, resulting in lower accuracy of small indel calls compared to NGS reads. When both small and large-structural variants need to be called, both NGS and TGS reads may be available. Integration of the two data types with unique error profiles could improve robustness of small variant calling in challenging cases. However, there isn’t currently such a method integrating both types of data. We present a novel method that integrates NGS and TGS reads to call small variants. We leverage the Mixture of Experts paradigm which uses an ensemble of Deep Neural Networks (DNN), each processing a different data type to make predictions. We present improvements in our DNN design compared to previous work such as sequence processing using one-dimensional convolutions instead of image processing using two-dimensional convolutions and an algorithm to efficiently process sites with many variant candidates, which help us reduce computations. Using our method to integrate Illumina and PacBio reads, we find a reduction in the number of erroneous small variant calls of up to ~30%, compared to the state-of-the-art using only Illumina data. We also find improvements in calling small indels in low-complexity regions.

## Introduction

Short-read sequencing or Next Generation Sequencing (NGS) technology from Illumina produces highly accurate reads of length typically between 50-150 bases. The high accuracy is an attractive aspect for calling variants - including substitutions and small indels (indel length ≤ 50bp). However, some inherent challenges remain, especially in calling small indels in low-complexity regions (Li, et al., 2018), where short reads may have difficulty unambiguously characterizing variations. In our experiments, we find that over 88% of erroneous indel calls made by a current state-of-the-art variant caller occur in the low-complexity regions of the genome.

Third Generation Sequencing (TGS) technology can produce sequencing reads that are thousands of bases long. Recent advances in PacBio Circular Consensus Sequencing (CCS) technology has improved the consensus base calling accuracy significantly. However, the remaining errors are predominantly small indels, and these errors are highly context-dependent and systematic, causing difficulties for a variant caller to reach the right consensus from multiple reads covering such locations. Hence, while TGS has been shown to call substitutions and large structural variants at high accuracy, the systematic errors cause small indel calling accuracy to be worse than that achieved using NGS reads (Wenger, Peluso, & Hunkapiller, 2019).

Both NGS and TGS reads have strengths and weaknesses, which make them suited for calling different variant types. NGS reads provide accurate small variant calls, while TGS reads provide accurate structural variant calls. Both types of variants may be relevant to different types of clinical applications such as in individualized medicine, for example, due to their role in influencing the individual’s susceptibility to diseases (Deng, Zhou, Fan, & Yuan, 2017) (Guo, et al., 2018). To call all types of variants, sequencing data from both sources should be considered and made available. Further, we speculate that if the sources of errors from the two platforms are in some sense complementary, it would be desirable to have a method that can take advantage of both data types to improve the accuracy of variant calling. Currently available methods for calling small variants can use only one of the two types of sequencing data exclusively, which is a limitation. Hence, using currently available tools, when both types of sequencing data are available, our only option is to call variants using one of the types of data exclusively.

Small variants are the most numerous types of variants in the genome. Both substitutions and small indels are widely associated with increased risk of many diseases such as different types of cancer (Stacey, et al., 2007) (Dai, et al., 2019) (Carlo, et al., 2020). Due to the complex nature of some of the associations, the role of variants is still being studied. Given that small variant callers today use only one type of sequencing data, and as a result consistently make erroneous calls in certain types of regions (e.g., indel calls in low-complexity regions) due to the error modes characteristic of a single sequencing technology, it is likely that the importance of variants in such regions may be less well-understood today. In addition, currently accepted benchmarks for variant calling such as Genome-In-A-Bottle (Zook, et al., 2019) have uncharacterized regions in the genome which may carry variants of significance. Some of these regions cannot be characterized due to the reliance, solely, on one type of sequencing data (namely short reads). Hence, a variant calling method using both short and long reads has the following potential advantages - (i) general reduction of errors in small variant calling, which is important from a clinical and research point of view, by leveraging long-read sequences that were sequenced for a different purpose (ii) reduction of errors in currently benchmarked, but challenging regions of the genome, improving our utilization of such regions in research and clinical settings, and (iii) enabling future studies into regions of the genome that are not currently benchmarked due to current attempts’ reliance on only one type of sequencing data.

Deep Neural Networks (DNNs) have recently become the preferred method for pattern recognition tasks and have been applied successfully to the problem of small variant calling (Poplin, et al., 2018). This was enabled by the availability of high-quality, benchmarked truth-sets from efforts such as the Genome-In-A-Bottle (GIAB) and Platinum Genomes (Eberle, et al., 2016). When designing a system with DNNs, the underlying patterns are discovered by the DNN from training data rather than expressed manually through models and algorithms by the method developer. DNNs can learn complex patterns from large amounts of data that would be infeasible for a method developer to examine manually to cover all underlying cases. In theory, DNNs can represent true underlying patterns in data due to their ability to operate as universal function approximators (Hornik, Stinchcombe, & White, 1989).

DNNs also provide mechanisms to combine data from multiple platforms. The canonical method to combine data for analysis is straight-forward. A sufficiently large neural network, in theory, is able to represent any function *f*(*I*, *P*) = *y*, based on its universal approximation ability, where *I*, *P* can be sequencing data from two different sequencing platforms and *y*, a set of variant calls. However, more sophisticated techniques can be designed which are more efficient in terms of computation, as well as more effective in terms of learning ability.

We have developed a caller, **H**ybrid **E**valuation of sma**LL** gen**O**mic variants (HELLO), which performs variant calling by integrating sequencing data from Illumina and PacBio reads. The method is based on the Mixture-of-Experts paradigm (Jordan & Jacobs, 1993) which allows multiple *experts* to make independent predictions, and a *switch*, or *meta-expert* to choose one of the experts’ predictions. Each expert in our implementation is itself a DNN specialized to the error profile of a different type of data (Illumina, PacBio, or both). The meta-expert is a DNN that learns what types of genomic regions are suited for each expert. In addition, each expert and meta-expert uses specialized architectures tailored for genome sequencing data, which allows us to reduce the computational complexity of the analysis. Applying our method to a whole-genome case-study for which both types of sequencing data are available, we find that it outperforms the state-of-the-art variant caller, that uses only Illumina reads. Our analysis also shows that the method improves accuracy of calling small indels in low-complexity regions of the genome, which are harder to call using either Illumina or PacBio reads, indicating that the effect of the errors in the individual sequencing platforms may be countered to some extent by HELLO due to each having different error characteristics.

## Results

### Methodology overview

**Figure 1.**
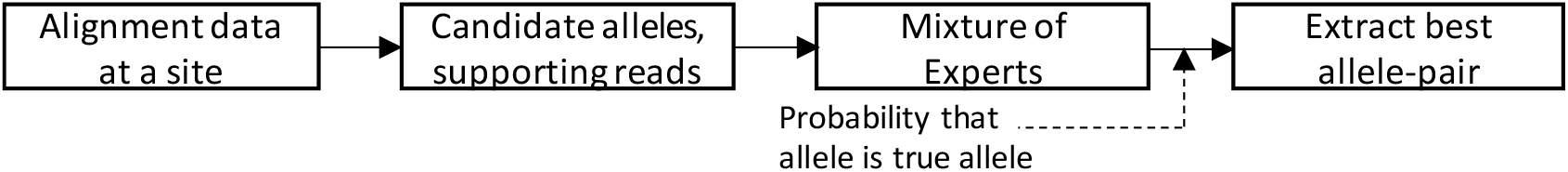
Using the MoE to predict variants. Candidate alleles and supporting reads are extracted at a site. The MoE uses this data to predict the probability that a candidate allele is a true allele at the given site. Every allele pair (non-unique combinations allowed) is then checked and the best pair is selected.

### Multi-label classification

In a typical germline paradigm, the human DNA consists of two haplotypes that may be represented as a single unique sequence of bases, or two different sequences of bases at any site in the genome; that is, there are one or two unique true alleles per site. Based on read alignment data at a given site, one may infer candidate alleles at the site. Some candidate alleles may arise from errors in sequencing and/or alignment. Hence, the number of candidate alleles and true alleles can vary from site to site. A variant caller needs to select one or two alleles as true alleles from a set of candidate alleles of variable cardinality. In machine learning terms, a model used for variant calling should be capable of labeling one or two categories as positive (as true alleles), from a variable number of categories (candidate alleles). This task differs from most popular DNN-based classification tasks (e.g., ImageNet image classification) which select a single positive category per input data, from among a fixed number of categories. Predicting multiple labels as opposed to a single label is called multi-label classification.

In our paper, we use the canonical method to solve multi-label classification; the independent prediction of the presence or absence of each category, otherwise called the binary-relevance method (R.Boutell, JieboLuo, XipengShen, & M.Brown, 2004). In terms of genome sequencing, this implies separately predicting the presence or absence of each candidate allele at a locus in the sequenced genome. The steps involved in predicting the ground-truth allele at a site are presented in Figure 1.

### Mixture of Experts

The core computations in our method are based on a DNN trained using the Mixture-of-Experts (MoE) framework for ensemble learning. In the MoE framework, there are multiple *experts* or sub-DNNs (henceforth simply referred to as “DNNs”), each providing predictions for a given input. A switch associated with a *meta-expert* then selects from the predictions of the multiple experts, based on a probability distribution assigned to the experts based on the input data by another DNN, which we refer to as the meta-expert DNN. During training we try to improve the likelihood of making the correct predictions using the MoE. This involves not only improving the predictive power of each expert, but also the predictive power of the meta-expert DNN, which essentially determines which expert is the most suited for a given input.

The MoE framework is attractive when considering prediction from multiple data sources with different error characteristics, and under the assumption that different sequencing technologies may work well for different regions in the genome. A straight-forward application of the MoE framework to our problem uses one DNN (expert) for each data-source.

Our actual implementation is slightly more nuanced, with the addition of a third expert as depicted in Figure 2. The third expert, NGS+TGS, is a monolithic DNN that accepts inputs from both NGS and TGS platforms and internally combines them into a single internal representation. Additional details regarding internal representations (features) and the architecture of the MoE is explained in the Supplementary Document. A single DNN operating on the combined representation has higher representational power compared to two individual DNNs independently processing one sequencing platform each. This may be reasoned as follows: the class of functions *f*(*A*, *B*) form a superset over the class of functions (*f*_1_(*A*), *f*_2_(*B*)). However, while the representational capacity may be higher, it may also be harder to train such a DNN. The MoE framework allows the NGS+TGS expert to focus on cases where the combination of the data from the two technologies is useful, with the option of using the individual experts for cases where learning becomes difficult using the combined data, but is simpler using one of the two types of data.

**Figure 2.**
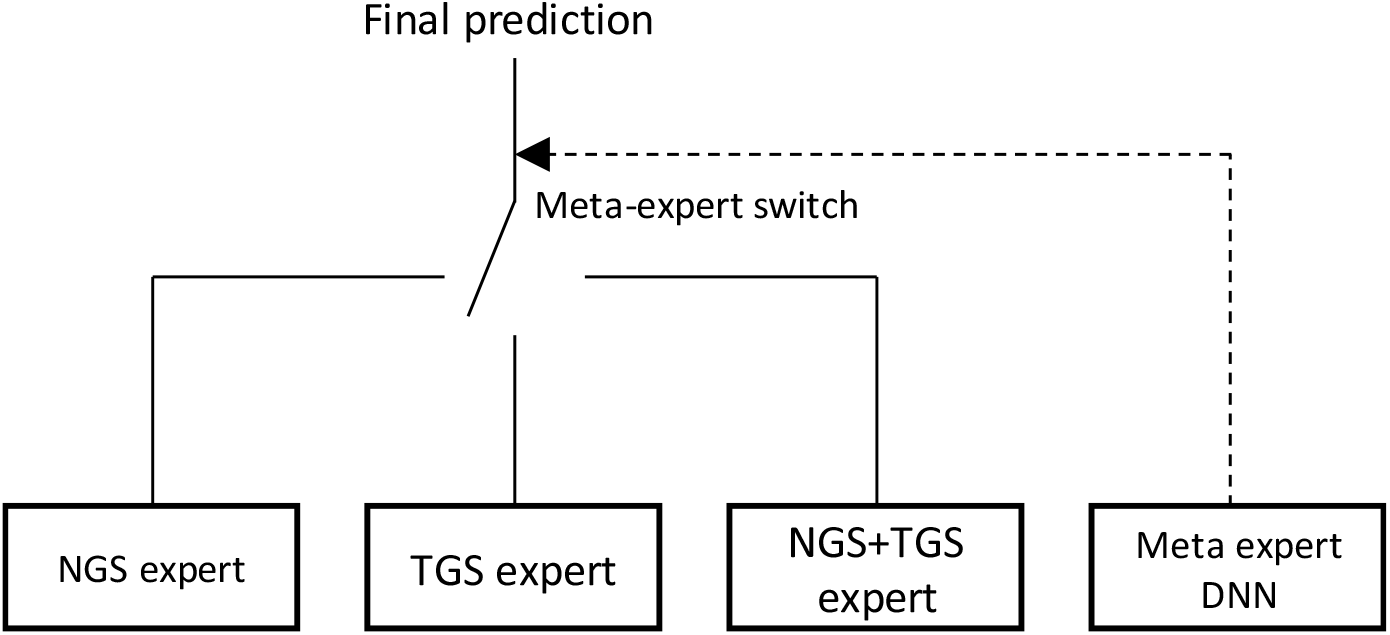
The MoE architecture of our model

### MoE architecture

Each Illumina and PacBio read is encoded into a multi-dimensional numerical array representing a sequence. Encoding of bases quality scores etc into numerical values, follows DeepVariant (Poplin, et al., 2018). When analyzing a site, these encoded read sequences are fed into the MoE, along with information on candidate alleles and their supporting reads.

The block diagram of the MoE is in Figure 3. The MoE uses one-dimensional (1D) Convolutional Neural Networks (CNN) which use residual connections (He, Zhang, Ren, & Sun, 2015). The MoE has different stages for processing read-level, allele-level and site-level information. The first stage is comprised of a pair of CNNs (“Read-level convolver” in Figure 3(a)-(b)) that process representations of either Illumina or PacBio reads. The outputs, which we refer to as read-level features, are sequences that emphasize the spatial information in each read relevant to variant calling. In the next stage, read-level features corresponding to a candidate allele are accumulated and passed through a second pair of CNNs (“Allele-level convolver” in Figure 3(a)-(b)) to obtain allele-level features summarizing evidence in favor of each candidate allele from each sequencing technology. Cumulation of allele-level features provide site-level features that summarize available data at a site, and the reference context. Combined allele-level and site-level features are also produced using additional CNNs (Figure 3(c)) which integrate evidence from both sequencing platforms to provide a single representation for allele-level evidence and site-level evidence. In the final stage, allele and site-level features are fed into four CNNs - three experts, and the meta-expert (Figure 3(d)). The NGS and TGS experts use allele and site-level features from the corresponding sequencing platforms, and the other two use the combined features. Internally, the experts compare allele-level evidence for each allele to the site-level evidence to determine whether the allele is noisy or not. The meta-expert uses the combined site-level feature to determine which expert it prefers at the site. We combine the outputs for the experts as follows.

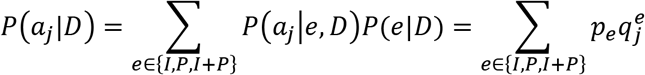

*D* represents the data at the site. *P*(*a*_*j*_|*D*) is the probability that allele *a*_*j*_ is a true allele according to the MoE, *P*(*a*_*j*_ |*e, D*) is the probability that *a*_*j*_ is a true allele according to the expert *e*. *P*(*e*|*D*) is the probability that expert *e* is selected by the meta-expert. Note that 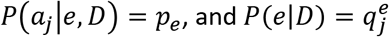, shown in Figure 3(d). We then proceed to determine the best pair of candidate alleles to perform the variant call at the site.

**Figure 3.**
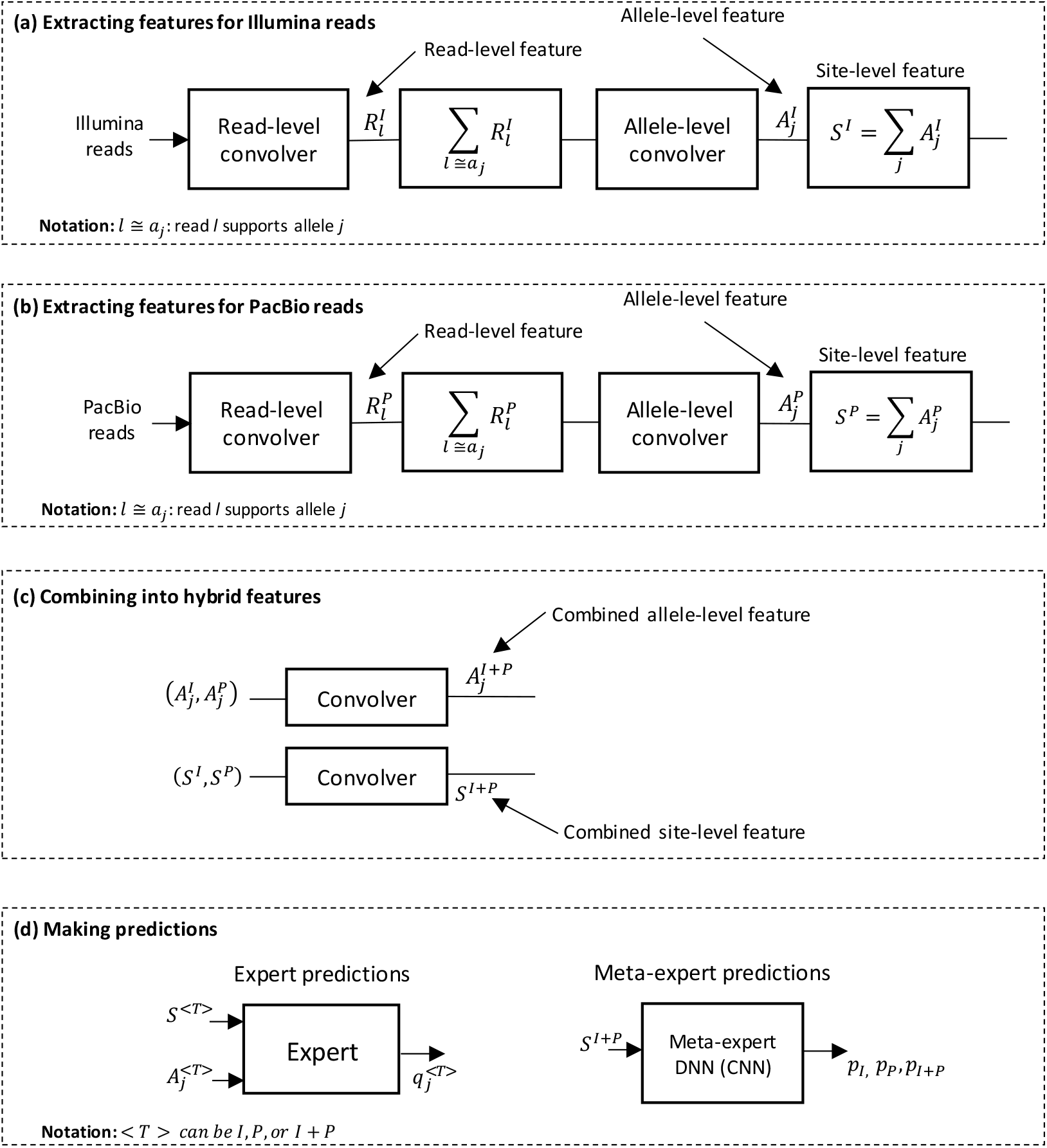
Block diagram of the MoE. NGS, TGS and NGS+TGS experts are indicated using the letters I, P, and I+P (I stands for Illumina, and P for PacBio). The term convolver in the figure refers to a CNN. (a)-(b) Generation of read-level, allele-level and site-level features for Illumina and PacBio data. (c) Combining features from two sequencing platforms. (d) Using allele and site-level features to make predictions; q^<T>^_j_ is the probability that allele j is a true allele according to expert <T>; p_<T>_ indicates the probability that the meta-expert chooses expert <T> at the given site.

While we follow the data encoding schemes introduced in DeepVariant, we present several architectural improvements in our DNN design. DeepVariant recycles an image processing DNN architecture, Inception v3 (Szegedy, Vanhoucke, Ioffe, Shlens, & Wojna, 2016), and converts sequencing data to fit the model’s original intended application. We recognize the nature of the data as composed of sequences. Hence, we do not use 2D convolutions that are used for pattern recognition in images, but 1D convolutions that are used for pattern recognition in sequences, which require less computations per layer than 2D convolutions. Convolutional layers in DeepVariant perform convolution operations not only along each read, but also across reads. Since reads may be assumed to be independently sequenced, and since DNNs operate well to capture structure in the data rather than random, independent events, we do not expect convolutions applied across reads to add useful operations and avoid them. Due to the design choices mentioned above, our model is comparatively small (< 6.5 million parameters) compared to DeepVariant (~25 million parameters). In addition, we do not use “null reads” that DeepVariant uses to pad input images to a fixed size, saving computations. DeepVariant performs predictions on allele pairs. This requires *O*(*n*^2^) DNN predictions per site when there are *n* candidate alleles at a site. Compared to this, we make predictions for each allele independently, requiring only *O*(*n*) predictions.

Detailed information on the architecture of the MoE, preparation of input data to the MoE, training, variant calling algorithm etc are provided in the Supplementary Document.

### Experimental validation

We used data from GIAB for the purposes of training our model, as well as testing its accuracy. Training data (including validation data for determining stopping condition of the training process; additional details are in the Supplementary Document) was compiled using alignment data from HG001, as well as ground-truth variants in benchmarked regions of the reference sequence as published in the GIAB repository. For benchmarking the accuracy of HELLO, we used HG002 whole genome data, which was not used in the training process. We compiled the training and benchmarking data as follows: 30x coverage Illumina data was prepared for HG001 and HG002 genomes. 30x and 15x data were compiled for PacBio Sequel II CCS sequencing technology for HG001 and HG002 genomes. HELLO models were trained on each of the combinations − 30x Illumina + 15x PacBio, and 30x Illumina + 30x PacBio using the HG001 dataset. For benchmarking the models, we used the corresponding combination from the HG002 genome. We prepared three variant call sets: two callsets from HELLO - one for 30x Illumina + 15x PacBio and one for 30x Illumina + 30x PacBio - as well as one callset from DeepVariant v0.9 with 30x Illumina sequencing data as input. The calls were evaluated using the hap.py evaluation tool (Krusche, et al., 2019).

In this paper, we focus on the numbers of mistakes—false positives and false negatives—made by the variant callers rather than the traditional metrics of Precision and Recall. Advances in variant calling have made success the common case rather than the exception, but work remains to perfect the solutions to the point that the results are more widely accepted in science and medicine. Traditional accuracy metrics show little change in response to reduction in errors for difficult-to-call sites, as these successes are masked by correct calls made at the vast majority of easier sites. Looking at the number of errors in callsets shifts the focus to those calls that are difficult to make. The numbers of errors made when calling small indels is summarized in Figure 4. Using PacBio and Illumina reads with HELLO reduces the number of FP errors by up to 27% and FN errors by up to 30%, compared to the results from DeepVariant. We further examine indel error characteristics within subsets of low-complexity regions in the genome which fall within benchmarked regions. Low-complexity regions are interesting regions to analyze. Majority of false positive indel calls from Illumina reads are found in low-complexity regions. Also, PacBio reads are susceptible to systematic indel errors in regions with repeat structures such as homopolymers which are found in low-complexity regions. It is interesting to see whether combining the two types of sequencing data, both susceptible to indel errors in low-complexity regions, but potentially due to different underlying causes and error models, allows us to improve the accuracy in these challenging regions.

**Figure 4.**
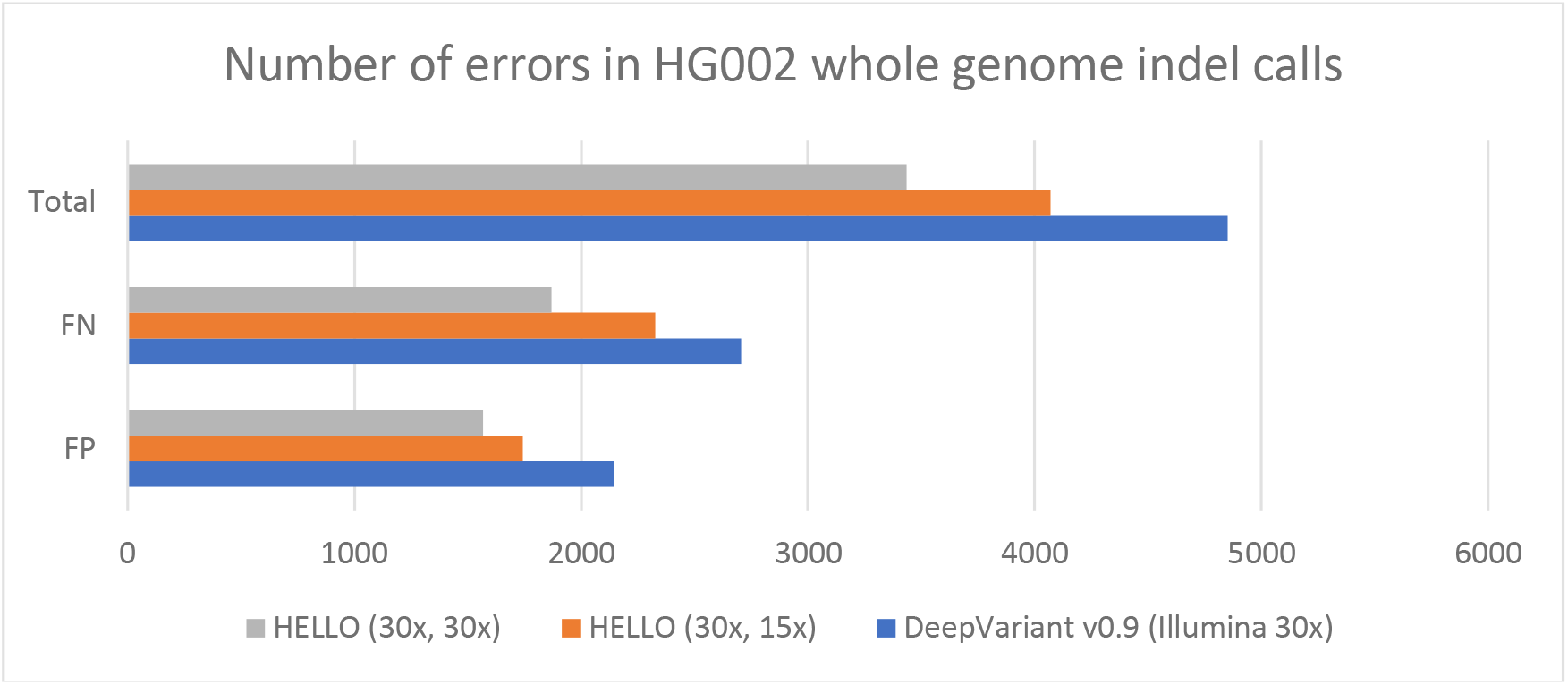
Number of Indel call errors (FP: False positives, FN: False Negatives). HELLO (Ax, Bx) implies HELLO using Ax coverage Illumina and Bx coverage PacBio. Notes: HELLO (30x, 30x) (Precision, Recall) = (0.9967, 0.9959); HELLO (30x, 15x) (Precision, Recall) = (0.9963, 0.9949); DeepVariant (Precision, Recall) = (0.9955, 0.9941)

The analyses are summarized in two sets of plots in Figure 5. Figure 5(a)-(c) use a set of low-complexity regions based on manually curated lists of collections of repeats in the reference sequence (Zook J., 2018). Figure 5(d) uses a second set of low-complexity regions as determined by the tool *sdust* (Morgulis, Gertz, Schäffer, & Agarwala, 2006) (Li, Minimap2: pairwise alignment for nucleotide sequences, 2018). *sdust* uses a method based on the abundance of triplets defined on the alphabet {A, C, G, T} in genomic regions, which allows assigning scores to the relative complexity of the region. Based on this score and a threshold, low-complexity regions may be identified. We use *sdust*-based low-complexity regions that were published previously (Li, et al., 2018).

**Figure 5.**
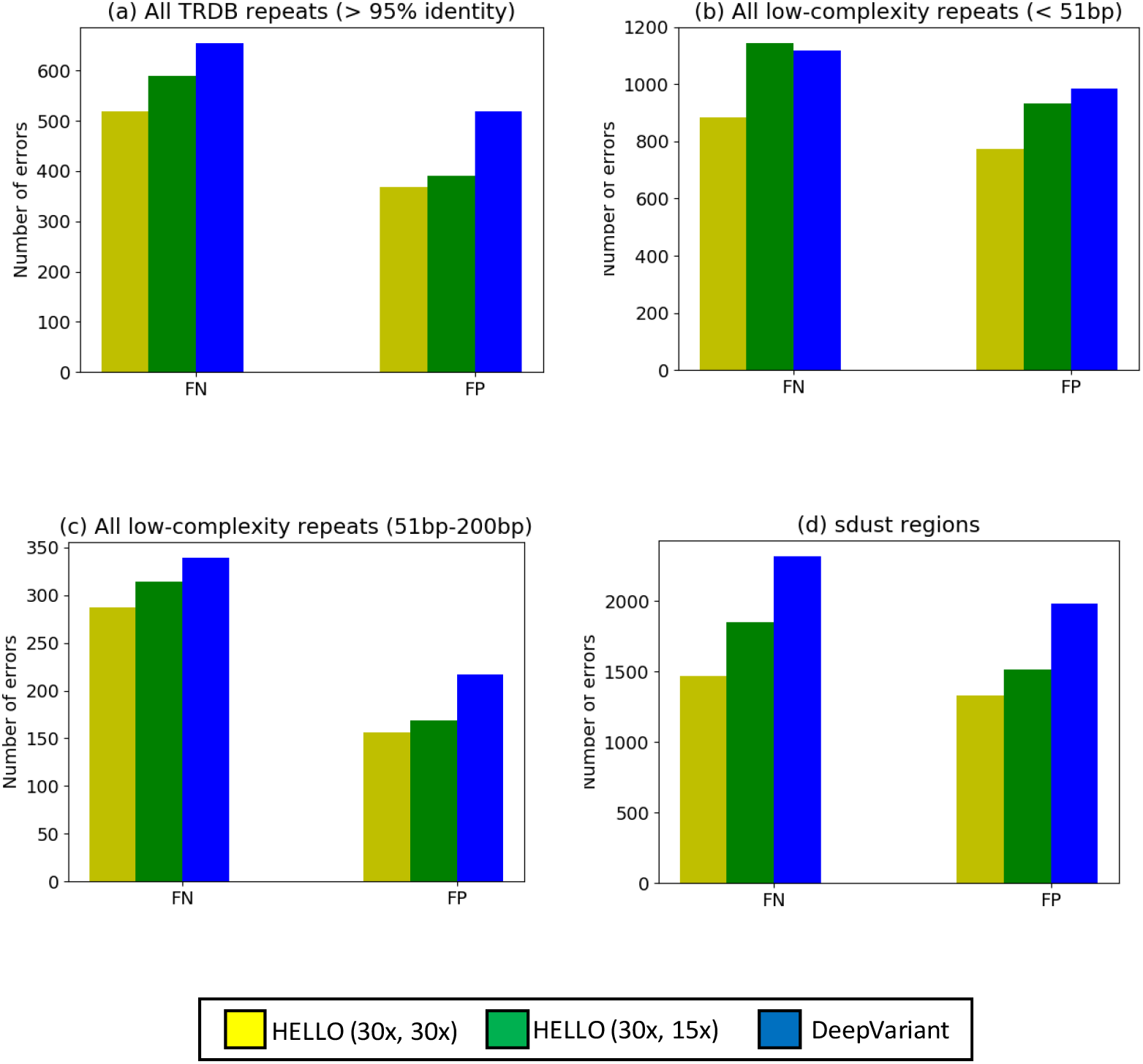
Stratification of indel call errors by region. Comparison between DeepVariant v0.9 using 30x Illumina reads and our method using configuration: Illumina 30x, PacBio 30x data, and configuration: Illumina 30x, PacBio 15x data

Regarding Figure 5(a)-(c), there are broadly two types of low-complexity regions that are split into three categories for analysis. The first type is a set of low-complexity regions summarized in the Tandem Repeats Database, TRDB, (Gelfand, Rodriguez, & Benson, 2006). This is a database of repeats in the human genome, out of which we examine repeats where the repetitive units have an identity of 95% (Figure 5(a)) or more. Some exact tandem repeats are not included in this database. The second set adds to the TRDB repeats mentioned above, a set of exact tandem repetitions as well. This set is further divided into two subsets - one where the repetitive region is small (Figure 5(b)), in other words, of the same size as the maximum number of inserted or deleted bases that is generally considered to constitute a *small indel* (< 51bp long), and one where the repetitive regions are longer, of lengths up to the order of the length of an Illumina read (Figure 5c; 51bp - 200bp). In all cases, our method improves false positive indel call accuracy compared to DeepVariant. Using 30x PacBio reads, both the number of false negative errors and the number of false positive errors are reduced in all three cases. Using only 15x PacBio reads, a similar result holds for TRDB repeats as well as low-complexity repeats 51bp-200bp long. In the case of shorter low-complexity repeats, the number of false negative errors is slightly worse and the number of false positive errors, better when compared to DeepVariant.

Unlike the manually curated low-complexity regions in Figure 5(a)-(c), the *sdust*-based regions in Figure 5(d) are a set of regions quantitatively determined to be low-complexity. Most of the erroneous calls in the stand-alone Illumina (88%), and HELLO (81% when using 30x Illumina, 30x PacBio) fall in low-complexity regions determined by *sdust*. While the manually curated low-complexity regions do capture a sizeable number of false calls, the *sdust*-based analysis captures many more false calls and encompasses almost all the errors. At the same time, *sdust*-based regions cover only ~2.1% of the complete reference, and ~1.6 % of the benchmarked regions in the reference. When looking at this exhaustive set of challenging regions in the genome, the use of PacBio reads reduces errors significantly compared to the other three categories. The improvement is more pronounced than in the case of manually curated repeat-based sets of low-complexity regions. This indicates that there are two types of low-complexity regions in the genome where Illumina reads are susceptible to errors, one where the context-dependent systematic errors in the PacBio reads result in only a modest improvement in accuracy, and one where such context-dependent errors are not as pronounced and where the improvement is more significant.

We next look at improvement in indel call accuracy with size of indel. This is presented in Figure 6. Three categories of indels are presented - indels of size 1-5bp, 6-15bp, and longer indels. The strongest improvement is observed for intermediate-sized indels (6-15bp). Illumina reads are likely to represent shorter indels more accurately than longer ones due to the relative size of short reads compared to indel size. Hence, as indel size increases, the signal strength for the indel also drops when using Illumina reads. We hypothesize that intermediate-sized indels present a sweet spot where combining Illumina and PacBio datasets sufficiently makes up for this loss of signal strength, even though PacBio reads have a higher indel error rate. Under this assumption, the NGS+TGS expert would be capable of making intermediate-sized indel calls with significantly higher accuracy than a caller that uses only one of the two types of sequencing data such as DeepVariant. If so, the meta-expert should be able to identify such scenarios and invoke the NGS+TGS expert in these cases. To see whether the NGS+TGS expert is indeed preferred in these cases, we examined 352 intermediate-sized indel errors in DeepVariant that were correctly called by HELLO (30x, 15x) (not all errors could be accurately tracked due to differences in indel variant representations among the different variant caller outputs and the evaluation program). We found that the meta-expert preferred the NGS+TGS expert in all but two of the cases, giving some support to our hypothesis.

**Figure 6.**
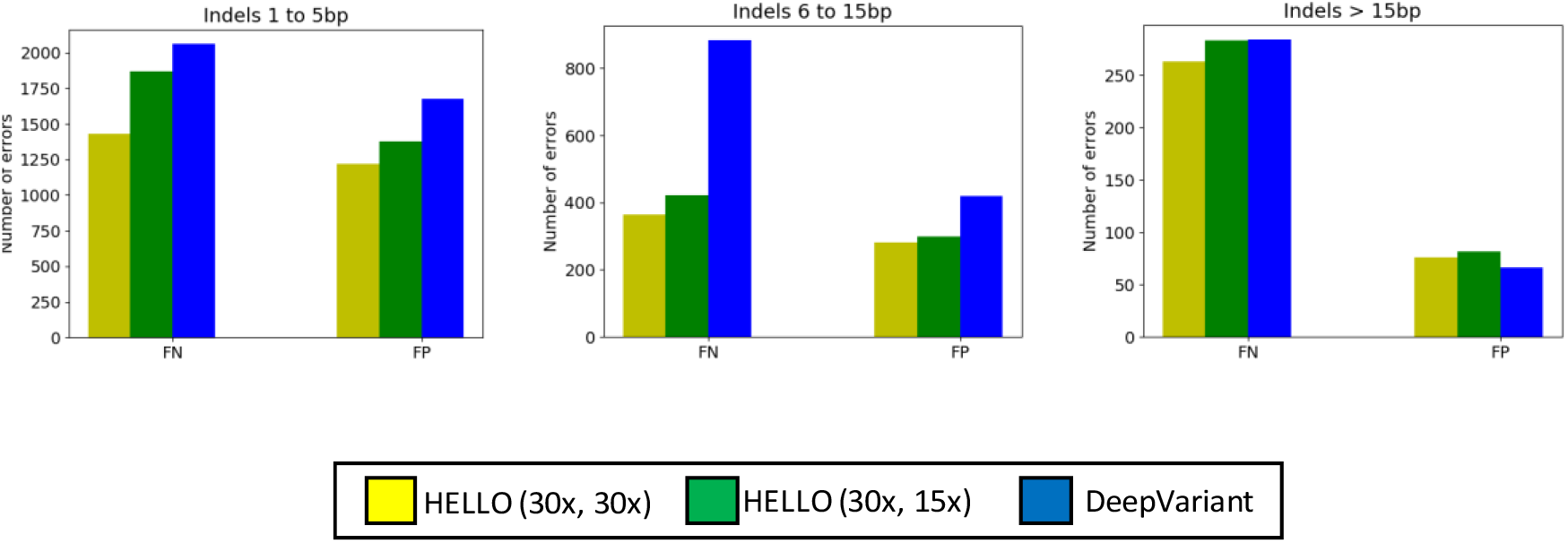
Analysis of indel errors by indel size

**Figure 7.**
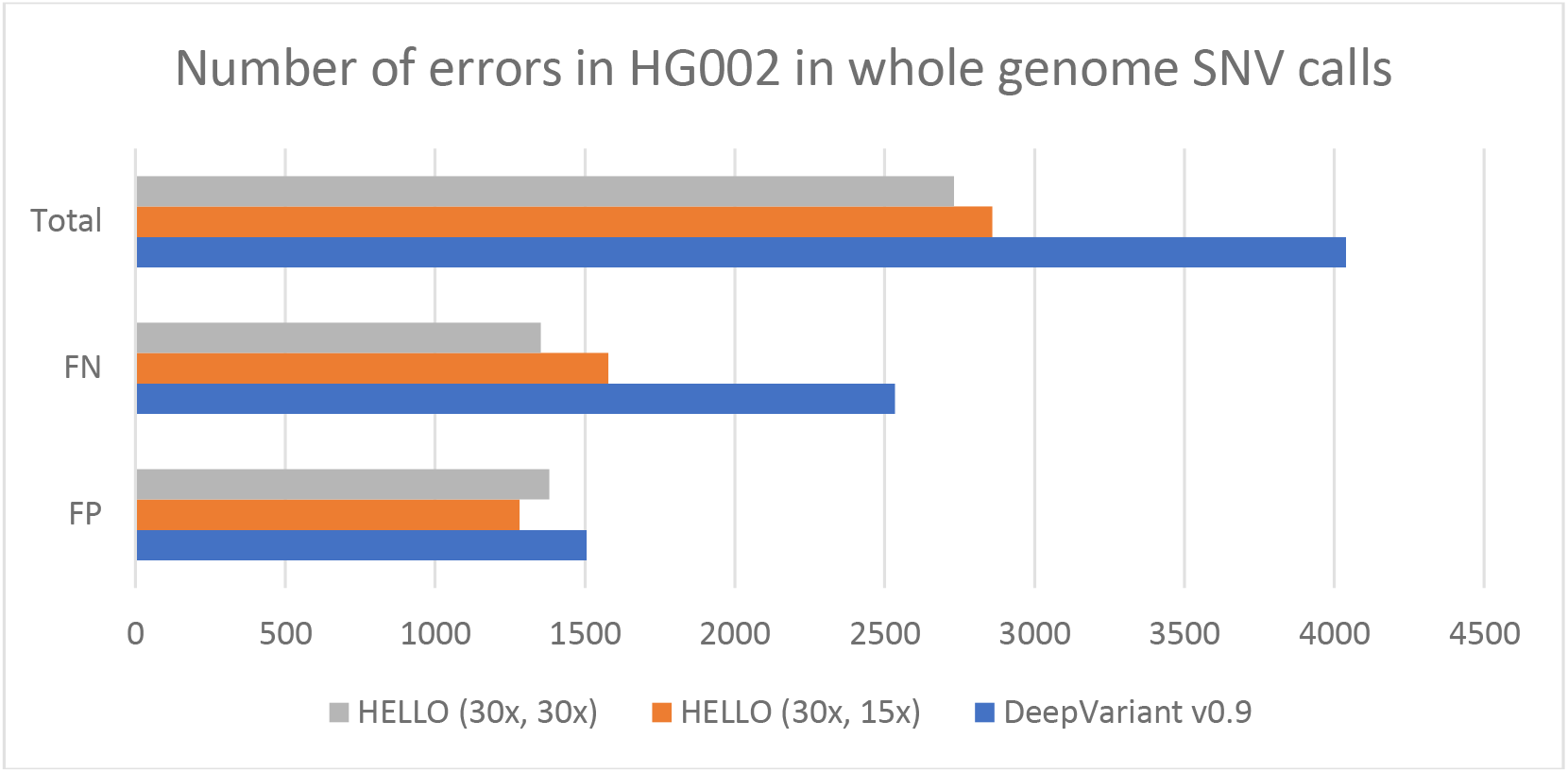
Number of errors when making SNV calls. Note: HELLO (30x, 30x) (Precision, Recall) = (0.9995, 0.9995); HELLO (30x, 15x) (Precision, Recall) = (0.9995, 0.9994); DeepVariant (Precision, Recall) = (0.9995, 0.9991)

Finally, we look at the number of errors made when calling substitution type errors. The total number of errors is reduced by up to 32% when using PacBio reads (30x Illumina, 30x PacBio). Using only 15x PacBio reads gives us 29% improvement indicating that a small amount of PacBio data can be helpful in cutting down errors significantly. Notably, the false negative error count is not significantly different from DeepVariant’s, though comparatively better, and significant improvements come from the reduction of false positive calls. Combining substitutions and small indels together, the net reduction in the number of erroneous calls is up to 30% (when using 30x Illumina and 30x PacBio reads).

## Discussion

We devised a hybrid variant calling method, HELLO, that combines PacBio and Illumina reads, data from sequencing platforms with different error profiles, but some common weaknesses. In cases where both types of sequencing data are available, using our method allows improving the accuracy of small variant calls over that of the state-of-the-art variant caller as seen in our experiments. In addition, we studied the performance of our method in calling small indels in contexts of low complexity in the reference, which has been historically difficult for small variant callers. Such contexts can cause problems for both Illumina reads and PacBio reads; almost all false positive indel calls in Illumina reads arise in these regions, and PacBio reads are susceptible to context-dependent indel errors in low-complexity regions. Our analysis indicates that using the hybrid method helps improve the accuracy in these regions, rather than compound the problem that both types of sequencing data have in these regions. Furthermore, within low-complexity regions, there are sub-regions where the gains are better or worse, indicating that in some cases the different error-profiles of the two sequencing platforms may be used to eliminate the effects, whereas in others it may not be as effective. Finally, when looking at improvements in indel calls with size of indel, HELLO provides outsized improvements for intermediate-sized indels (6-15bp long) pointing to some fundamental advantage in integrating NGS and TGS data. We examined many of the errors made by DeepVariant in this category that were correctly called by HELLO and traced the calls to the NGS+TGS expert indicating that hybridizing the data may be the key to calling these indels with high accuracy.

## Availability

- Source-code is released open source at https://github.com/anands-repo/hello
- Experimental data is available online. Please refer to links provided in the source-code repository. Details on data preparation can be found in the Supplementary Document.

## Supplementary information

### Determining hotspot regions and candidates

We iterate through reads from an alignment file counting the number of reference-matching and reference-deviating cigar strings (reference-deviating cigars include substitutions and indels). If the number of reads supporting a reference-deviating cigar at a location is high enough to warrant a closer look, we mark the location as a hotspot. This is similar to the method used in DeepVariant.

During variant call, reads aligning to a window surrounding a contiguous set of hotspots are extracted and the hotspot analysis is performed again in the region with slightly higher sensitivity. Adjacent hotspots triggered by indels are coalesced into a single site of interest. This minimizes the amount of conflict in the variants reported. The remaining hotspots are individually treated as sites of interest. The sites of interest are analyzed to determine candidate alleles and supporting reads, which are fed into the Mixture of Experts (MoE) in HELLO. A read supports a candidate allele if it contains the candidate allele aligned to the site of interest. If a read partially overlaps a site of interest, then we see whether the partial overlap may be resolved by unambiguously matching it to one of the other candidate alleles at the site. If so, the read is determined to be supporting that candidate. If not, the read is discarded.

### Details of input data representation, and read filtration

**Figure 1.**
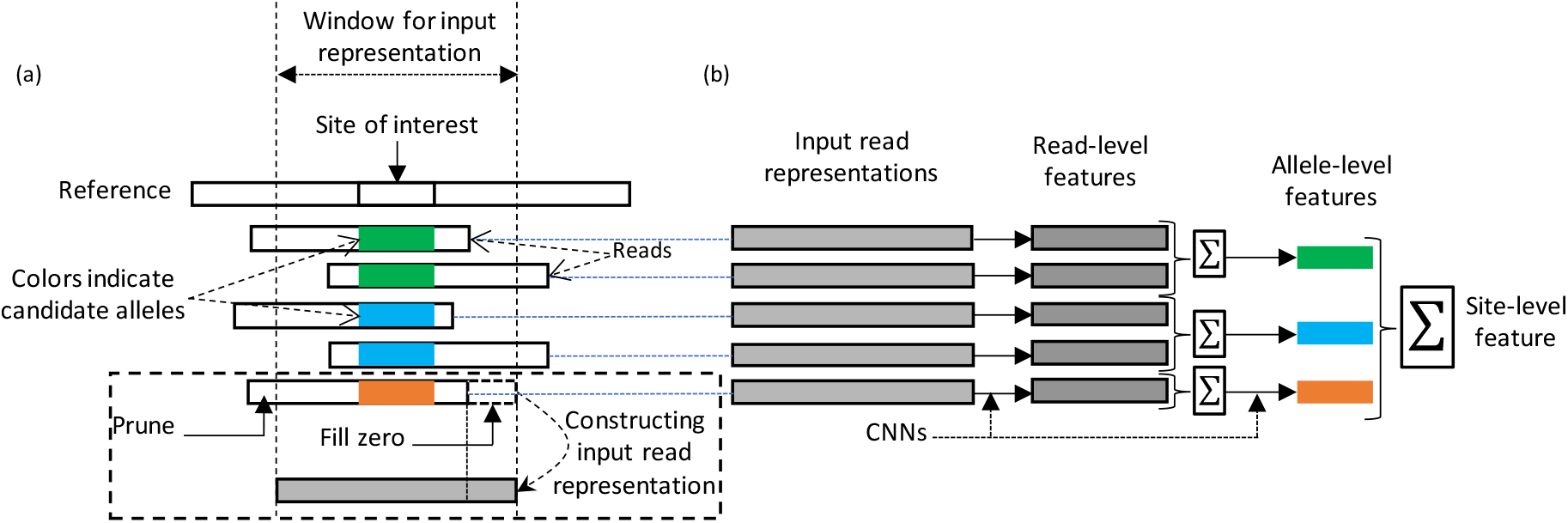
(a) Alignment data at a site of interest, and conversion of reads into representations for the DNN. The portion of a read within a window around a site of interest is extracted and the bases, quality scores etc are converted to a sequence of vectors. Zero vectors are padded to this sequence corresponding to parts of the window that are unoccupied by the read, if the read doesn’t fully occupy the window. (b) How input read representations are converted to different features for a single sequencing platform (NGS or TGS). The input representations pass through multiple stages giving different features summarizing read-level, allele-level and site-level evidence.

**Figure 2.**
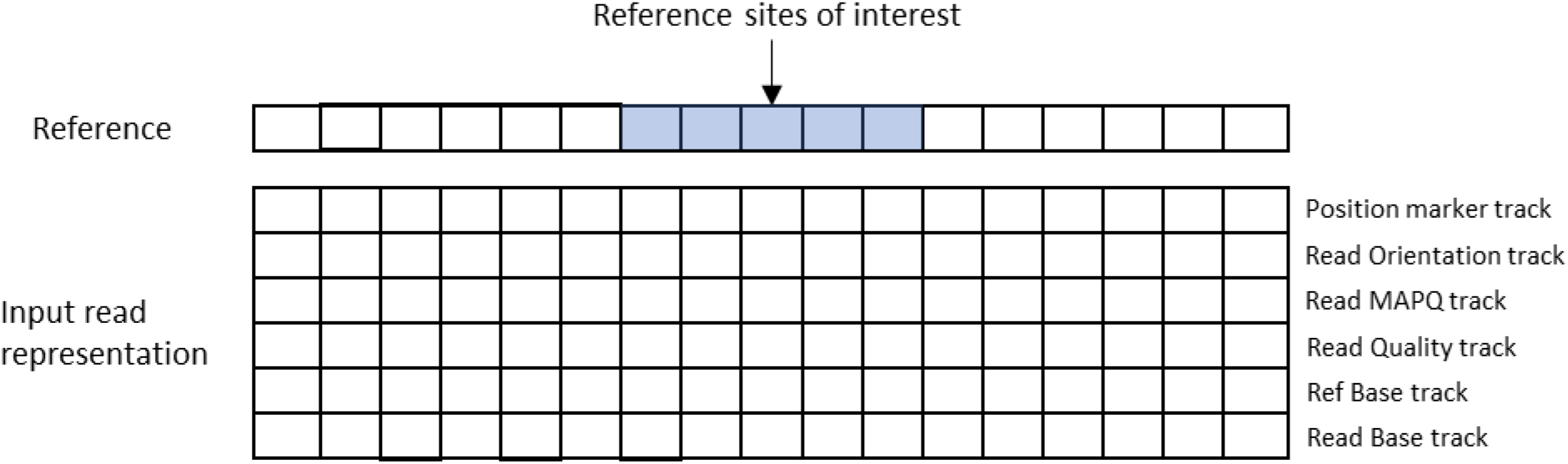
Details of input read representation. The input read representation is a six-dimensional vector, each dimension encoding some characteristic of the read, its alignment, or the reference base it aligns to

Figure 1 (a) shows a putative site carrying a variant that we want to analyze using the MoE. At the site, there are a certain number of candidate alleles, and a certain number of reads supporting each candidate allele. These numbers vary from site to site. The Mixture of Experts (MoE) accepts input data at the granularity of individual reads. Each read aligning to the site of interest is extracted and converted to a sequence representation. The sequence represents a fixed-size window from left to right, around the site of interest. Each element in the sequence is a six-dimensional vector representing a position within the window. The dimensions are the read base (A, C, G, or T), the reference base it aligns to, base quality, mapping quality of the read, forward/reverse strand and whether the element corresponds to a position within the site of interest, or whether it is a surrounding contextual location. A read may only occupy a portion of this window in which case, zeros are filled in for the unoccupied portion. Portions of the read that lie outside the window are discarded.

Details regarding the input read representation are illustrated in Figure 2. The representation method takes cues from DeepVariant. The following are the similarities with DeepVariant.

1. The representation contains values from 0 to 255 which may be represented using a single byte. This allows us to store the data with a lower storage footprint and provides a nice dynamic range.
2. A, C, G, T are represented using non-zero levels within 0 to 255 (in the Read Base track) and are nicely distributed in that range. Insertions are represented with a zero at the base just before the insertion. Deletions are represented using zeros for all deleted bases as well as the base just before the deletion.
3. Quality scores and Mapping quality scores are capped and scaled to fit evenly within the 0 to 255 range (Read quality track and Read MAPQ track). Quality, *qual*, is converted as 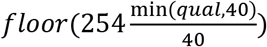, Mapping quality, *MAPQ*, is converted as 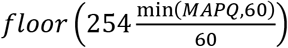.
4. Read orientation (forward/reverse strand) is encoded and applied to the Read orientation track. Two values, 70, 240 are used to encode the read orientation.

The following are differences compared to DeepVariant:

1. Individual reads are encoded as separate sequences. In DeepVariant, reads aligning to a single location are encoded into 2D images.
2. Instead of simply allotting one of the tracks to saying whether a read base at that position matches or mismatches with respect to the reference, the actual reference bases are encoded here (Ref base track; base encoding is similar to that of read bases).
3. We also mark the site of interest in the image, that is, the locus at which we are testing the presence of a variant. This is encoded in the position marker track in the image above. Sites of interest are marked using a value 240, and contextual positions are marked with a value 70.

The length of the read representation used is 150, that is encoding a total of 150 reference positions. A read that doesn’t align to the complete 150 base positions is filled with zeros in positions the read doesn’t span. A read that spans beyond the boundary of the 150 bases is truncated. Hence an input read representation represents a 6-dimensional sequence of length 150.

Read filtration also follows criteria similar to that used in DeepVariant. Following are the criteria:

- Reads with mapping quality < 10 are not used
- Reads which span the reference sites of interest with one or more low quality bases (< 10) are not used
- Reads which are not mapped in a proper pair (when paired-end reads are used) are discarded
- Supplementary alignments and duplicates are discarded
- Reads failing vendor quality checks are discarded

### DNN design

The MoE within HELLO is designed to have multiple stages. The data at the boundary of each stage may be considered to represent information at different levels - arranged as read-level, allele-level and site-level. Hence, as information flows through the MoE, higher and higher-level information is composed and consumed. The data representations at the boundaries of the stages are in the form of multi-dimensional arrays emphasizing aspects of the data important for variant calling as may be interpreted by subsequent stages in the model. They are referred to as “features” in the sequel. A schematic of the data representations and the general flow of feature creation in the DNN is presented in Figure 1(b). Below we describe the features in the MoE and how they are generated/processed. The block-diagram of the MoE is shown in Figure 3 (reproduced from the Main Document). The complete diagram of the MoE is given later in this document.

**Figure 3.**
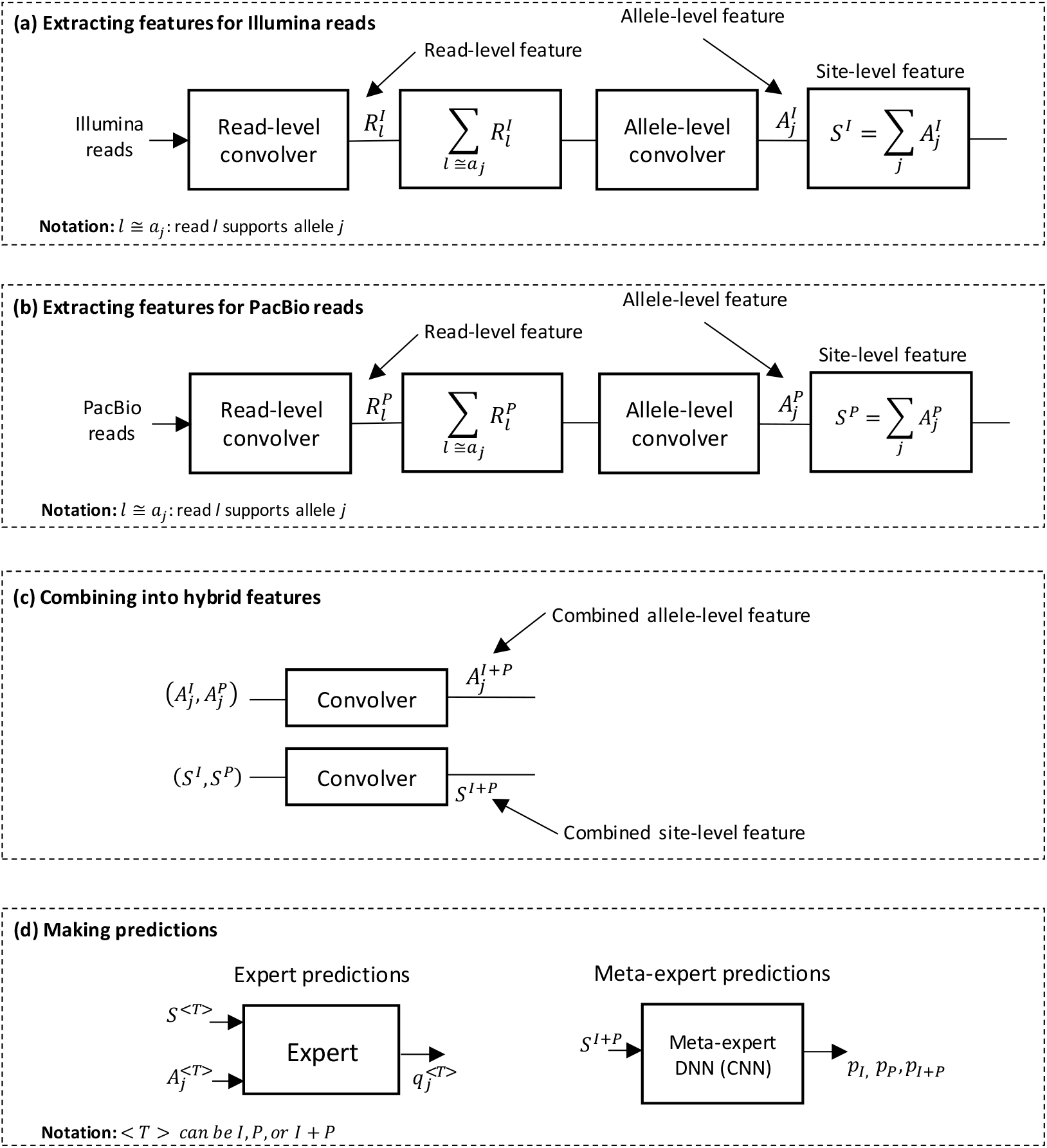
Block diagram of the complete MoE. NGS, TGS and NGS+TGS experts are indicated using the letters I, P, and I+P (I stands for Illumina, and P for PacBio).

#### Read-level features

A Convolutional Neural Network (CNN), labeled the “read convolver” in Figure 3(a)-(b), processes the input read representations. CNNs can extract temporal patterns in the read sequences and encode them into the output representation. The output of the read convolver represents read-level features. We use separate read convolvers to process the data from each sequencing platform to obtain two sets of read-level features at each site, so that each CNN is acclimatized to the error-profiles of the sequencing platform it is handling.

#### Allele-level features

While the read-level features summarize a single read, allele-level features incorporate information from all the reads that support an allele. There are two steps performed in producing the allele-level features - i) adding the read-level features of reads that support an allele, and ii) passing the result through a CNN (“allele-level convolver” in Figure 3(a)-(b)). These steps are performed separately for read-level features from the two sequencing platforms to obtain two sets of allele-level features. We assume that read-level features are additive, in the sense that, adding read-level features provides a meaningful feature for analysis by the allele-level convolver. While the raw input encoding of the reads does not represent such *superposable* features, read-level features are produced as the result of processing the input using the read-level CNN. Following the Deep Learning paradigm, we assume that there exist functions which produce outputs that are superposable, and that the read-level convolvers learn such functions during training.

The description above applies to allele-level features for a single sequencing platform. The MoE also involves DNNs that use allele-level features from both sequencing platforms - namely the NGS+TGS expert and the meta-expert DNN. The features for use by these DNNs are produced by combining allele-level features from the two sequencing platforms using CNNs (Figure 3(c)).

#### Site-level features

We assume that allele-level features may be superposed, like read-level features. Allele-level features at a site are added together to form site-level features. Site-level features may be considered to summarize the reference context of the site, as well as the type and amount of data available at a site. Like the case of allele-level features, we use a CNN to combine site-level features from two different sequencing platforms.

The allele- and site-level features are used to make predictions using the experts and the meta-expert (Figure 3(d)). The experts and the meta-expert DNN are implemented using CNNs. To determine the presence or absence of a candidate allele at the site of interest, each expert compares the allele-level feature for the candidate allele to the site-level feature at the site. This allows the expert to compare the evidence supporting the candidate allele for which a prediction is made, to the remaining evidence at the site and determine whether there is significant signal supporting the candidate, or whether it is the result of noise. The meta-expert uses the combined site-level feature to determine which expert is better suited to predict labels for the given input data. The combined site-level feature summarizes the characteristics of the site and the amount of data available from each technology to facilitate such a prediction.

### MoE operation during training and calling

The loss function for training the Mixture of Experts (MoE) is derived using the Expectation Maximization (EM) algorithm. To invoke the EM algorithm MoE is considered a generative model of labels when the inputs are given. Under this assumption, the meta-expert’s decisions become a latent, or hidden variable. The EM algorithm is used to solve problems where a generative latent variable model is to be fitted to some observations, and the latent variables’ values for each data point are unknown. The EM algorithm involves the Expectation step (E-step) where the expectation of the joint log likelihood of the observations and the latent variables is computed over the posterior probability of the latent variable. This gives us a cost function. This cost function is maximized in the next step, which is called the Maximization step (M-step). In our case, we perform this maximization through stochastic mini-batch gradient descend.

The cost function for the MoE from the E-step is given by

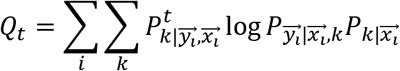

where *Q*_*t*_ is the cost function computed for the *t*^th^ training step, *k* is the identity of the expert (for three experts *k* takes three values), *y*_*i*_ is the output label and *x*_*i*_ is the input tensor data for the *i*^*th*^ training example. 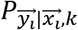 represents the probability of the label given the input data and assuming that the meta-expert chooses the expert *k* (in other words, the probability of the output label according to the *k*^th^ expert). 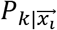 represents probability that the meta-expert chooses the *k^th^* expert (this is simply the *k*-th output of the meta-expert). 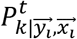 represents the posterior probability of the meta-expert’s choice being *k* given the input tensor, and its label. This can be computed easily from the DNN outputs through Baye’s rule. The superscript *t* indicates that this term is a constant, in that, it is a numerical value that doesn’t add terms to gradient computations for the MoE parameters; it is computed based on the DNN parameter values as of the *t*^th^ training step. This is in keeping with the requirement that this is the probability distribution over which the expectation (in the E-step) is calculated. The other two probability terms build backpropagation graphs and lead to parameter updates during the M-step.

During variant calling, given the predictions from the experts and the meta-expert, we need to combine them into a single prediction per allele. This results in the predictions of the experts being weighted by the probability distribution determined by the meta-expert DNN. This will be explained with simplified notation. For a given site and candidate allele (allele *j*, say), assume that the meta-expert DNN predicted the probabilities *p*_*I*_, *p*_*P*_, *p*_*I+P*_ for the experts, and the predictions from the experts for the candidate allele are 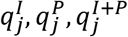. The quantities 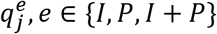 indicate the probability that candidate allele *j* is a true allele at the site according to expert *e*. Then, the mean prediction from the MoE for allele *j* is

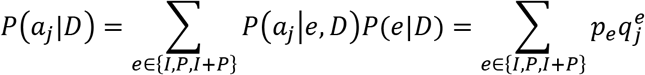

Given that there are *n* candidate alleles at a site, and *P*(*a*_*j*_ |*D*), or *P*(*a*_*j*_) for short, being the probability of allele *j* being present at the given site as predicted by the MoE, then the predicted allele pair *i, j* for the site is determined using the following equation; here 1_{.}_ is the indicator function.

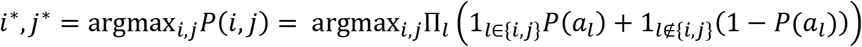

Note that the equation may be modified for any ploidy without modifying the architecture of the DNN itself, since the DNN is simply predicting the presence or absence of individual alleles. Hence, with minor changes, our method may be applied to genomes with different ploidies.

To determine the confidence of the prediction based on the best allele pair, we simply substitute *i*^∗^, *j*^∗^ into the expression for *P(i,j)*.

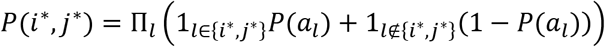

In these expressions, for alleles that are predicted to be at the site we use the probability of the allele being present, and for the alleles which have not been predicted to be at the site, we use the probability of the allele being absent, as predicted by the MoE. The above equations are probabilistically valid since each allele is predicted independently of the rest.

### Preparing training data, and performing variant calling

HELLO accepts two BAM files - one for Illumina reads and one for PacBio reads.

The BAM file for Illumina reads is pre-processed using GATK to determine consensus indels and realign the reads accordingly. This is a necessary step for short reads because read aligners align reads independently. If the read doesn’t cleanly span an indel location with enough flanking bases on both sides of the indel, the indel may not appear in the alignment. For example, independent alignment of reads can result in artifacts such as soft clipping of valid bases or reads may not all represent the indel identically. The problem is exacerbated in areas containing indels adjacent to short tandem repeats where the read needs to span not only the indel, but also the accompanying tandem repeat region in order to capture the correct indel. The local assembly-based procedure implemented in GATK can correct for such disagreements among short reads by determining consensus haplotypes in the region and aligning the reads to the consensus haplotypes. In the case of long reads, such issues are less pronounced since almost all reads covering an indel location span the entire indel location with enough flanking bases.

The pre-processed BAM files are analyzed to determine sites of interest. Candidate alleles are extracted from bases aligned to the sites of interest, reads supporting each candidate allele are determined, and representations are created for the reads.

To create training examples, we also require a set of ground-truth variants, as well as a set of benchmarked regions of the genome where the ground-truth variants set is valid. Hotspot locations outside of the benchmarked regions are discarded, and locations within benchmarked regions are labeled using the ground-truth variants set and a haplotyping algorithm. The labels and the read representations are written to disk.

During variant calling, the read representations (without labels), are fed into the trained MoE instance to determine prediction probabilities for each candidate allele. Once we have the probabilities for the presence of all candidate alleles at the site, we can determine the probability of every pair of candidate alleles at the site using the simple rules of probability (we are interested in pairs of candidate alleles because the human genome is diploid). Then the allele-pair with the best likelihood is selected for the site. The predicted likelihood for the selected allele-pair is converted to a variant quality score and then printed out in the VCF format.

### Additional details regarding implementation/training

The implementation of the MoE is in pytorch (version 1.2). For training the two models, 30x15x (30x Illumina, 15x PacBio), and 30x30x (30x Illumina, 30x PacBio), we tried two different learning rates, 0.001, and 0.002 each. The training was performed using data from human chromosomes 1 to 20 in the HG001 genome. Early stopping was used to terminate training. For 30x15x, we terminated training after two consecutive epochs gave no improvement in validation loss (validation loss is computed using random data-points from among chromosomes 1 to 20). For 30x30x, we terminated training after three consecutive epochs gave no improvement in validation loss. Variant calling was performed on Chromosomes 21, and 22 in HG001 to select the better model for each case. The selected models for 30x15x and 30x30x used learning rates 0.001, and 0.002 respectively. For training purposes, two different types of hardware platforms were available to us. One type used the IBM Power 8 architecture with 4 NVIDIA K80 GPU cards. The other type had an x86-based architecture with 2 NVIDIA V100 GPUs.

### Datasets

Links for downloading alignment data

HG001 Illumina: ftp://ftp-trace.ncbi.nlm.nih.gov/giab/ftp/data/NA12878/NIST_NA12878_HG001_HiSeq_300x/NHGRI_Illumina300X_novoalign_bams

HG001 PacBio: ftp://ftp-trace.ncbi.nlm.nih.gov/giab/ftp/data/NA12878/PacBio_SequelII_CCS_11kb

HG002 Illumina: ftp://ftp-trace.ncbi.nlm.nih.gov/giab/ftp/data/AshkenazimTrio/HG002_NA24385_son/NIST_HiSeq_HG002_Homogeneity-10953946/NHGRI_Illumina300X_AJtrio_novoalign_bams

HG002 PacBio: ftp://ftp-trace.ncbi.nlm.nih.gov/giab/ftp/data/AshkenazimTrio/HG002_NA24385_son/PacBio_SequelII_CCS_11kb

The reference sequence is hs37d5.

samtools (v1.9) view command (option -s) was used for creating the datasets used in experiments from the downloaded bam files.

Additional pre-processing of these BAM files (populate Read Groups field for the reads as required by GATK), was performed using Picard tools (version 2.21.1 @ https://github.com/broadinstitute/picard/releases/download/2.21.1/picard.jar).

To perform indel realignment on the Illumina data we used GATK Queue version 3.8-1.

The docker image of DeepVariant version 0.9.0 was used to run DeepVariant.

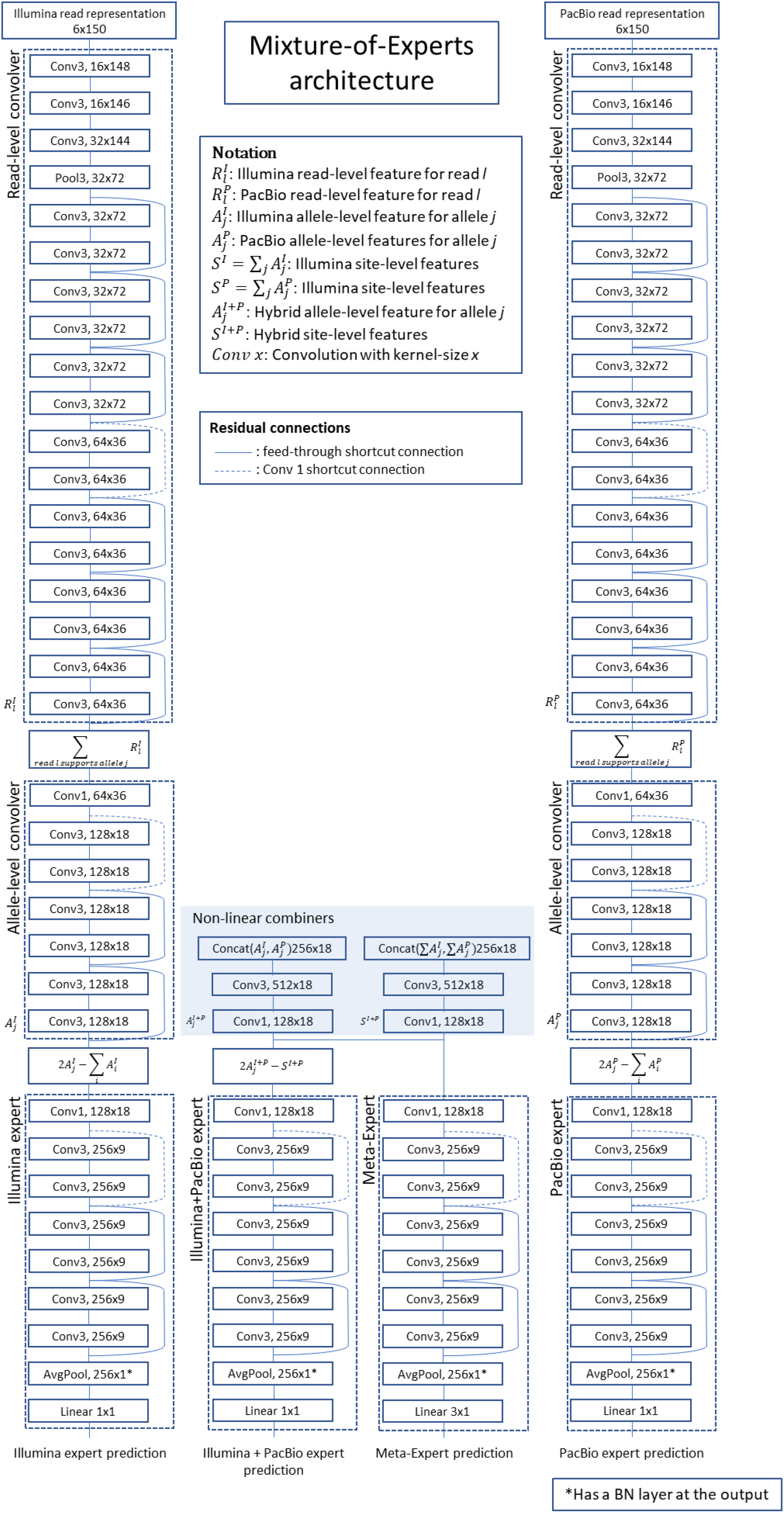

Note that each Conv layer in the figure has a convolutional kernel, followed by Batch Normalization and a ReLU activation function. As may be seen, experts and the meta-expert involve 34 or 36 neural network layers (excluding the Batch Normalization and ReLU activation functions). Non-linear combiners marked in the figure accept allele-level or site-level features from the two different sequencing platforms and combine them into a single feature with a non-linear function parameterized using a two-layer convolutional neural network.

### Other comparisons

The latest docker image for GATK was downloaded on 01/28/2020. BQSR, HaplotypeCaller, CNN2d scoring and FilterVariantTranches were run. GATK resources files used are as follows

- dbSNP for BQSR

- ftp://ftp.ncbi.nih.gov/snp/organisms/human_9606_b151_GRCh37p13/VCF/All_20180423.vcf.gz
- Resource files for variant filtration

- ftp://ftp.broadinstitute.org/bundle/hg19/hapmap_3.3.hg19.sites.vcf.gz
- ftp://ftp.broadinstitute.org/bundle/hg19/Mills_and_1000G_gold_standard.indels.hg19.sites.vcf.gz

Number of indel call errors in HG002 whole genome call is reported below.

| Method | FN calls | FP calls |
| --- | --- | --- |
| GATK – PASS (filtered calls) | 5432 | 3210 |
| GATK – All (unfiltered) | 4810 | 3374 |
| DeepVariant | 2705 | 2147 |
| HELLO (30x, 15x) | 2327 | 1743 |
| HELLO (30x, 30x) | 1869 | 1566 |

Number of SNV call errors in HG002 whole genome call is reported below.

| Method | FN calls | FP calls |
| --- | --- | --- |
| GATK – PASS (filtered calls) | 5837 | 2504 |
| GATK – All (unfiltered) | 5127 | 4507 |
| DeepVariant | 2535 | 1505 |
| HELLO (30x, 15x) | 1578 | 1281 |
| HELLO (30x, 30x) | 1352 | 1380 |

## Acknowledgements

This material is based upon work supported by the National Science Foundation (NSF) under Grant Nos. CNS 1624790, and CNS 1337732. Any opinions, findings, and conclusions or recommendations expressed in this material are those of the author(s) and do not necessarily reflect the views of the National Science Foundation.

